# A novel universal algorithm for filament network tracing and cytoskeleton analysis

**DOI:** 10.1101/2021.01.04.425230

**Authors:** D.A.D. Flormann, M. Schu, E. Terriac, D. Thalla, L. Kainka, M. Koch, A.K.B. Gad, F. Lautenschläger

## Abstract

The rapid development of advanced microscopy techniques over recent decades has significantly increased the quality of imaging and our understanding of subcellular structures, such as the organization of the filaments of the cytoskeleton using fluorescence and electron microscopy. However, these recent improvements in imaging techniques have not been matched by similar development of techniques for computational analysis of the images of filament networks that can now be obtained. Hence, for a wide range of applications, reliable computational analysis of such two-dimensional (2D) methods remains challenging. Here, we present a new algorithm for tracing of filament networks. This software can extract many important parameters from grayscale images of filament networks, including the Mesh Hole Size, and Filament Length and Connectivity (also known as Coordination Number. In addition, the method allows sub-networks to be distinguished in 2D images using intensity thresholding. We show that the algorithm can be used to analyze images of cytoskeleton networks obtained using different advanced microscopy methods. We have thus developed a new improved method for computational analysis of 2D images of filamentous networks that has wide applications for existing imaging techniques. The algorithm is available as open-source software.

## 1. Introduction

In recent decades, investigation of the filaments of the cytoskeleton, such as actin, vimentin, and microtubules, has become increasingly important for our understanding of cellular functions^1–4^. For example, the spatial organization of the cytoskeletal network has an important role in cell migration^5, 6^, cancer metastasis^7, 8^, and cellular mechanics^1, 9–12^. The methods used to detect and image these structures vary from low-resolution fluorescence imaging, through high-resolution fluorescent imaging, to electron microscopy^3, 6, 13–16^. The resolution of conventional light microscopy allows imaging down to 200 nm, and super-resolution microscopy can now detect features with a resolution down to around 20 nm. However, for the full structural networks, the resolution down to the atomic scale of electron microscopy is required.

The structures in these images can then be analyzed computationally, and the most commonly used method to investigate networks of grayscale imaging techniques are based on segmentation or ‘skeletonization’, such as with ImageJ plugins; e.g., DiameterJ and NeuronJ^17, 18^. These methods are user friendly and have well-defined and convenient output parameters. However, as the user has to define the ‘best segmented image’ prior to the analysis, these methods rely upon the subjective analysis of the user. This thus reduces the reliability and reproducibility of these methods. More reliable analysis can be provided by vectorial-based algorithms that allow more objective analysis of filaments in grayscale images^2, 19^. However, a limitation of this method is that the output parameters are commonly dedicated to a specific interest, and can therefore be limited.

To facilitate our ongoing analyses of scanning electron microscopy images of actin microfilament networks in cells, we have developed an algorithm that can analyze a wide range of filamentous networks that are imaged using different techniques, such as fluorescence microscopy, electron microscopy, and commercial photography techniques. We have named this algorithm the filament network-tracing algorithm (FiNTA). To the best of our knowledge, the FiNTA analyses more parameters than other computational tools designed for network analysis. These parameters include the Filament Length, the Connectivity (also known as the Coordination Number, and the Persistence Length in combination with the Angle Distribution. Thus, this algorithm can be used to analyze any kind of network in grayscale two-dimensional (2D) images.

## 2. Materials and methods

### 2.1. Vectorial filament network-tracing algorithm

The FiNTA is based on vectorial tracing of grayscale images. Therefore, although binary images can be analyzed, there is no need to create binary images prior to an analysis. To recognize the filaments in grayscale images, the FiNTA first identifies the directions of the filaments using image convolution with the Hessian matrix of the Gaussian kernels; thus, by validation of every pixel in terms of their intensities^20^. This is followed by the generation and connection of nodes along the filaments, which results in the tracing lines (Fig. 1A). It is possible to manually adjust the sensitivity of the method to any grayscale image by changing the input parameters, as described in Supporting Information (Table S1). For a more detailed description of the working principles and the mechanisms behind the algorithm, please see Supporting Information. The FiNTA identifies the filaments in a suitable time range; depending on the network complexity, the time range can be from seconds to minutes. It then automatically extracts the important network parameters. The algorithm is available on: https://github.com/SRaent/Actin.

**FIGURE 1.**
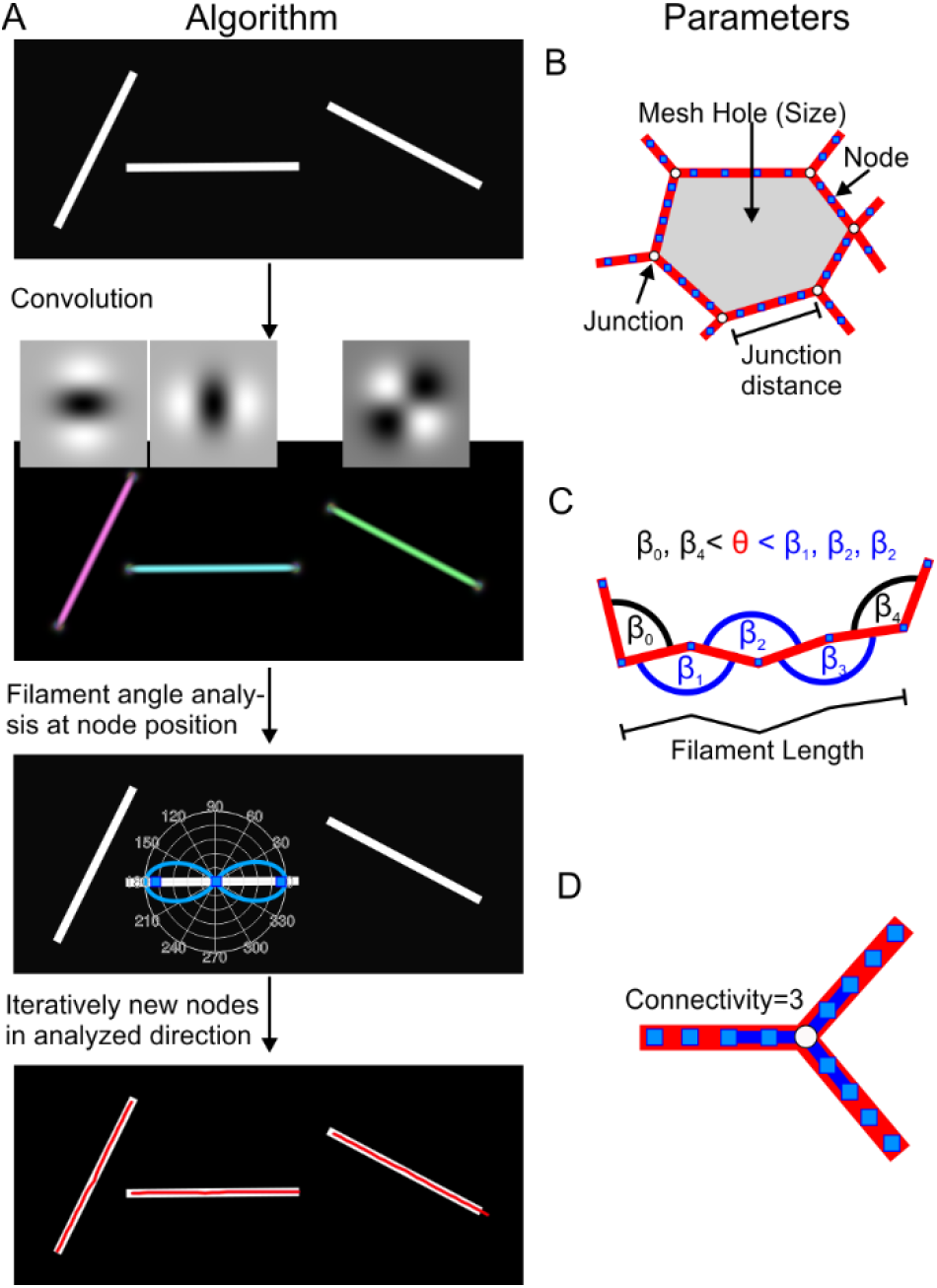
Graphical illustration of the four parameters implemented in the filament network-tracing algorithm (FiNTA). (**A**) Algorithm routine to trace filaments vectorially. Filament thickness: 12 px. (**B**) A single Mesh Hole with several junctions. (**C**) The Filament Length FL calculation. The break-off angle is θ (or smaller). The break-off angle assigned to the FiNTA is 180°-θ (as for all other angles), or larger. (**D**) Implementation of the Connectivity summarizes the nodes along the unification distance shown as the thick blue line. In the example, four nodes (blue squares) per fiber are connected to identify one filament as belonging to the junction (white circle).

### 2.2. Extracted parameters

The FiNTA provides information on at least eight of the most relevant parameters, which are: Mesh Hole Size; Circularity of each mesh hole; Junction Distance; Filament Density, Filament Length, Connectivity of each unified Junction, Global Angle Distribution, and the Persistence Length.

The Mesh Hole Size is the measure of the network pore size, pore area, or hole size, or similar, and this is defined as the area of the pores or holes within the network (Fig. 1B). The Circularity is defined as Ci = 4*π*(*MHS*)/*P*^2^, where *P* is the perimeter of a Mesh Hole. Consequently, the Circularity is a measure of the fractal dimension of the mesh holes. The Filament Density is the total network length divided by the total Mesh Hole Area. Furthermore, the Junction Distance as the length of a filament between two junctions (Fig. 1B). The Filament Length is the distance between nodes before reaching a defined break-off angle θ within two adjacent connections (Fig. 1C). The number of filaments that are connected to a junction is described with the term Connectivity, which in three-dimensional (3D) data analysis is often referred to as the Coordination Number. As a value of two represents a single filament, the minimum value of the Connectivity is three, while the maximum value is unlimited. For instance, in actin networks, the Connectivity rarely exceeds six, and these filaments typically have a mean connectivity of around 3.4. A user-defined unification distance summarizes the nodes to one junction (Fig. 1D). The Global Angle Distribution Distribution is defined as the angles of the connections between the nodes relative to the image orientation, where the left and bottom edges of the image represent the y-axis and x-axis, respectively. The worm-like chain model approximation is used to extract the Persistence length^21^.

To demonstrate which parameters are easier to extract with the FiNTA, we analyzed three scanning electron microscopy images of structurally different actin microfilament networks in cells. These were taken over the nucleus, in the perinuclear area, and in a lamellipodium at the cell edge, as shown in Figure 2A. We thereby determined the density and structure of the actin microfilament network, and how it differed between the different subcellular regions, according to 18 different parameters (Fig. 2B).

**FIGURE 2.**
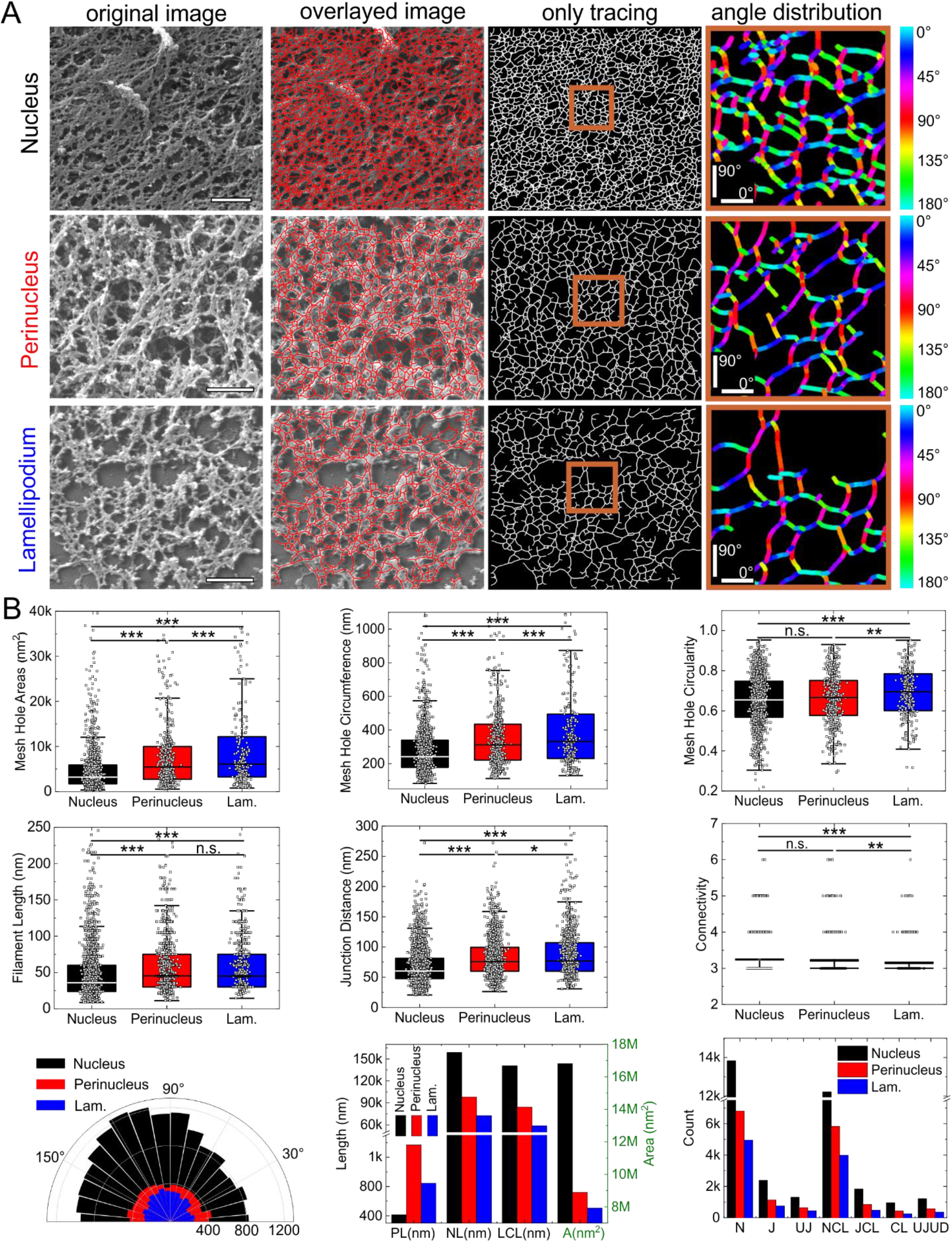
Scanning electron microscopy examples and parameters. (**A**) Representative actin images showing the tracing and angle distributions of three different cell regions of RPE1 cells. Scale bars and edge length of angle distribution images: 500 nm. (**B**) Quantification of 18 parameters extracted by the FiNTA. Lam., lamellipodium; PL, persistence length; NL, total network length (to calculate, e.g., Fiber Density LCL, total network length of closed loops; A, total Area of all loops; N, total number of nodes; J, total number of Junctions; UJ, total number of United Junctions upon user defined Unification Distance used to calculate the Connectivity; NCL, total number of Nodes that contribute to Closed Loops; JCL, total number of Junctions that contribute to Closed Loops; CL, total number of Closed Loops; and UJUD, total number of United Junctions of the network that only contain closed loops with the user-defined Unification Distance.

### 2.3. Biological sample preparations and imaging

The preparation of the biological samples and the imaging details are in the Supporting Information.

## 3. Results

### 3.1. Filament density, Filament Angles and Mesh Hole Size

The filament density is commonly calculated by determining the length of the total traced network, and dividing by the image size. To quantify the quality of this analysis, a realistic range of values of filament packing/densities were tracked that were within an acceptable error (10% SD). An orthogonal grid of white filaments on a black background was digitally created. In this network, the number of filaments was then increased, from a filament density of 92% white and 8% black pixels, to 26% white and 74% black pixels, as shown in Figure 3A-D. The Sizes and Counts of the Mesh Holes in these images were then analyzed using the FiNTA, which were compared to the known Sizes and Counts of the designed Mesh Holes. The results with the FiNTA were similar to the known Mesh Hole Sizes and Counts (Fig. 3E, F). This indicated that the FiNTA provided an accurate description of these porous and dense networks, with detection of up to 92% of the filaments.

**FIGURE 3.**
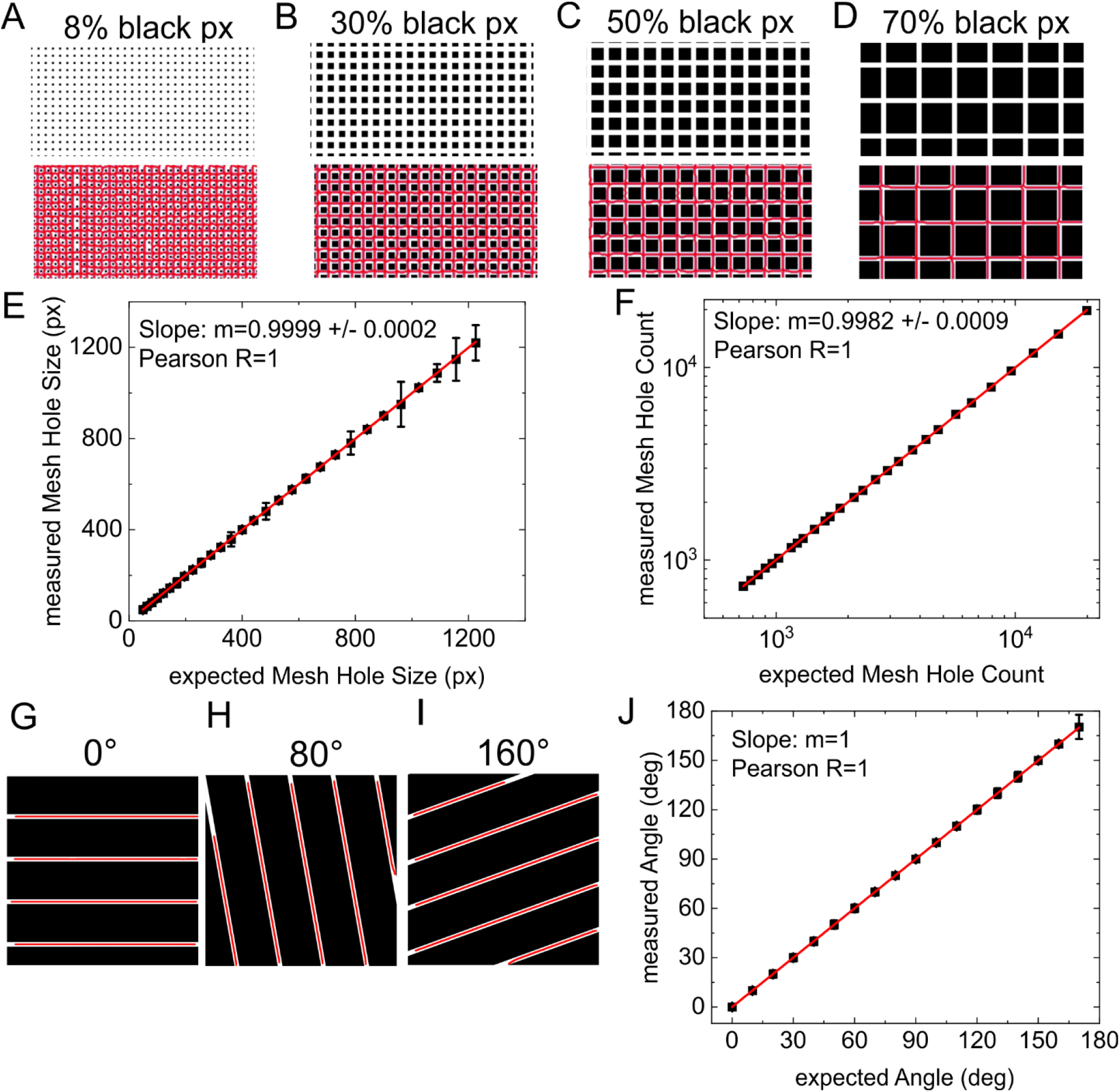
Fiber density and orientation quality tests. (**A-D**) Cropped regions of exemplary images with increasing black pixel fractions (decreasing filament density), with tracing results in the bottom half. Line thickness: 6 px. (**E, F**) Linear approximations of expected *versus* measured Mesh Hole Size (E) and Mesh Hole Count (F). (**G-I**) Exemplary images of different global Filament Angles with a line thickness of 27 px, that lead to accurate tracing lines. (**J**) Linear approximation of expected *versus* measured Angles.

To quantify the accuracy of the FiNTA analysis of the global Filament Angles, 18 images were created with Filament Angles from 0° to 170°, in 10° increments (Fig. 3G-I). The following FiNTA analysis of the Filaments Angles was very similar to the expected values, with a Pearson’s correlation R = 1 (Fig. 3J). This indicated that the FiNTA provided an accurate analysis of the Global Filament Angles.

### 3.2. Image resolution and noise level

Image resolution and the signal-to-noise ratio govern the quality of the various imaging methods. We thus determined how these features of images influence the FiNTA tracing of networks. For this, test images were initially created with a filament density grid of 74% black. To recognize different network types, the FiNTA detection of different grid sizes was also tested by analysis of lines of six different sizes of pixels.

For the signal-to-noise ratio, six digital noise levels were added, as infinite to zero signal-to-noise ratios, and the noise levels were calculated. The tracing quality of the FiNTA was then analyzed at these different signal-to-noise ratios by comparisons of the measured to the expected Mesh Hole Size and Count with the similarities expressed as percentage errors (Fig. 4). Here, the higher the resolution of the image, the more noise was acceptable, and *vice versa*. Therefore, the FiNTA can trace images across a wide range of image resolution, and with various noise levels.

**FIGURE 4.**
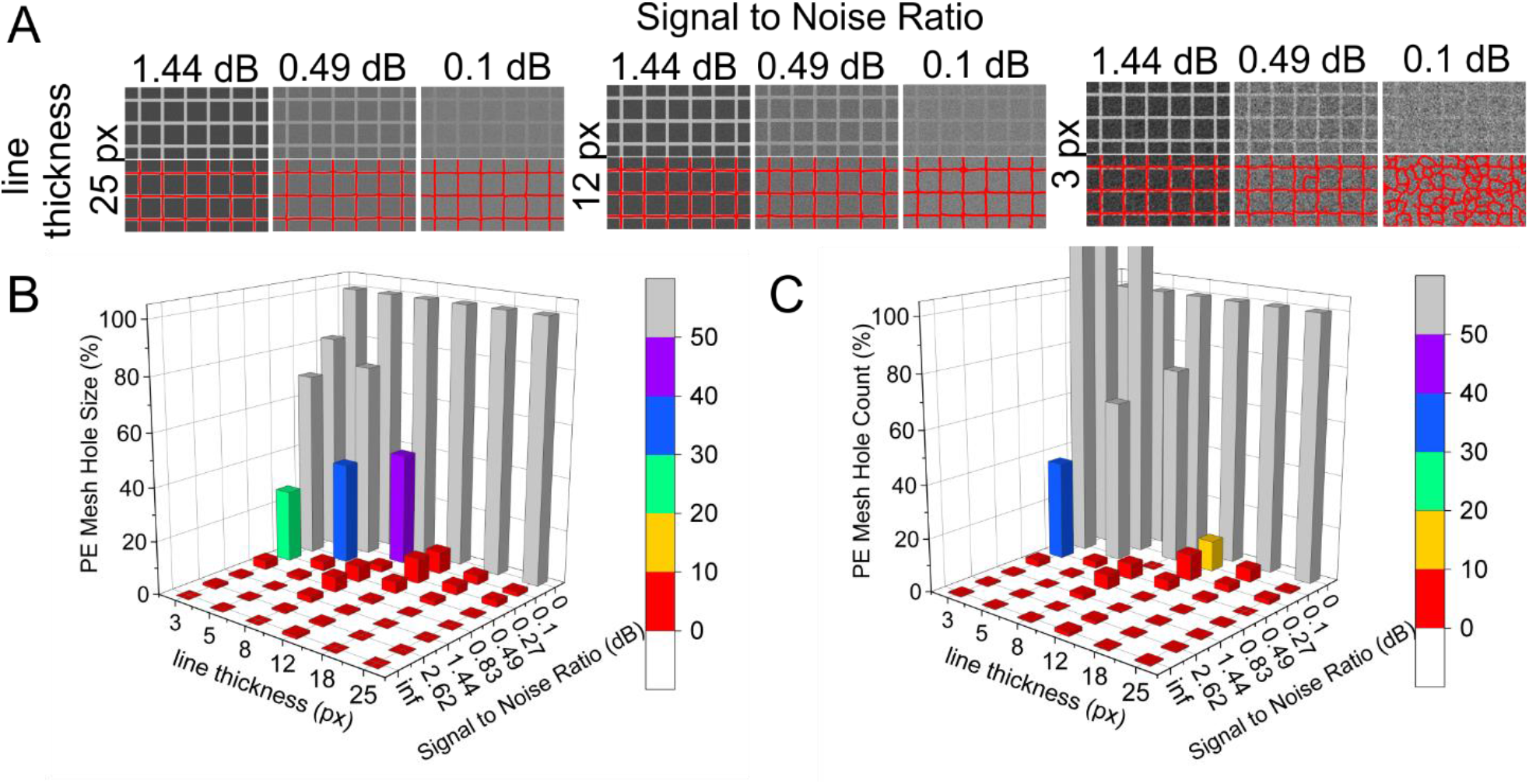
Mesh Hole Size by varying signal-to-noise ratio and line thickness. (**A**) Exemplary images of traced Mesh Holes by varying signal-to-noise ratio and line thickness. (**B**) Percentage errors (PE) of expected Mesh Hole Size and Count *versus* measured Mesh Hole Size (B) and Count (C), with dependence on signal-to-noise ration and line thickness.

The typical signal-to-noise ratios of fluorescence and electron microscopy images of the cellular actin cortex are in the range of 5 dB to 15 dB.

### 3.3. Connectivity

The Connectivity (also known as Coordination Number or Branching Number) is rarely included in algorithms for segmentation-based network analysis, and to the best of our knowledge, it has not been included in vectorial tracing algorithms. Nonetheless, it is an essential parameter that is important to include and describe in any comprehensive analyses of networks. Therefore, the Connectivity is also determined in the FiNTA.

To determine the efficiency of the Connectivity measured by the FiNTA, four black and white 2D test images were artificially created where the only Connectivity values were three, four, five, and six. As a Connectivity of two represents a line, the minimum value of the Connectivity is three.

The Connectivity measured on the original images with the FiNTA was then compared to the expected values, as shown in Figure 5A-D. These FiNTA values matched those expected (Fig. 5E, F), and therefore, the FiNTA can measure all of the predominant Connectivities within 2D networks.

**FIGURE 5.**
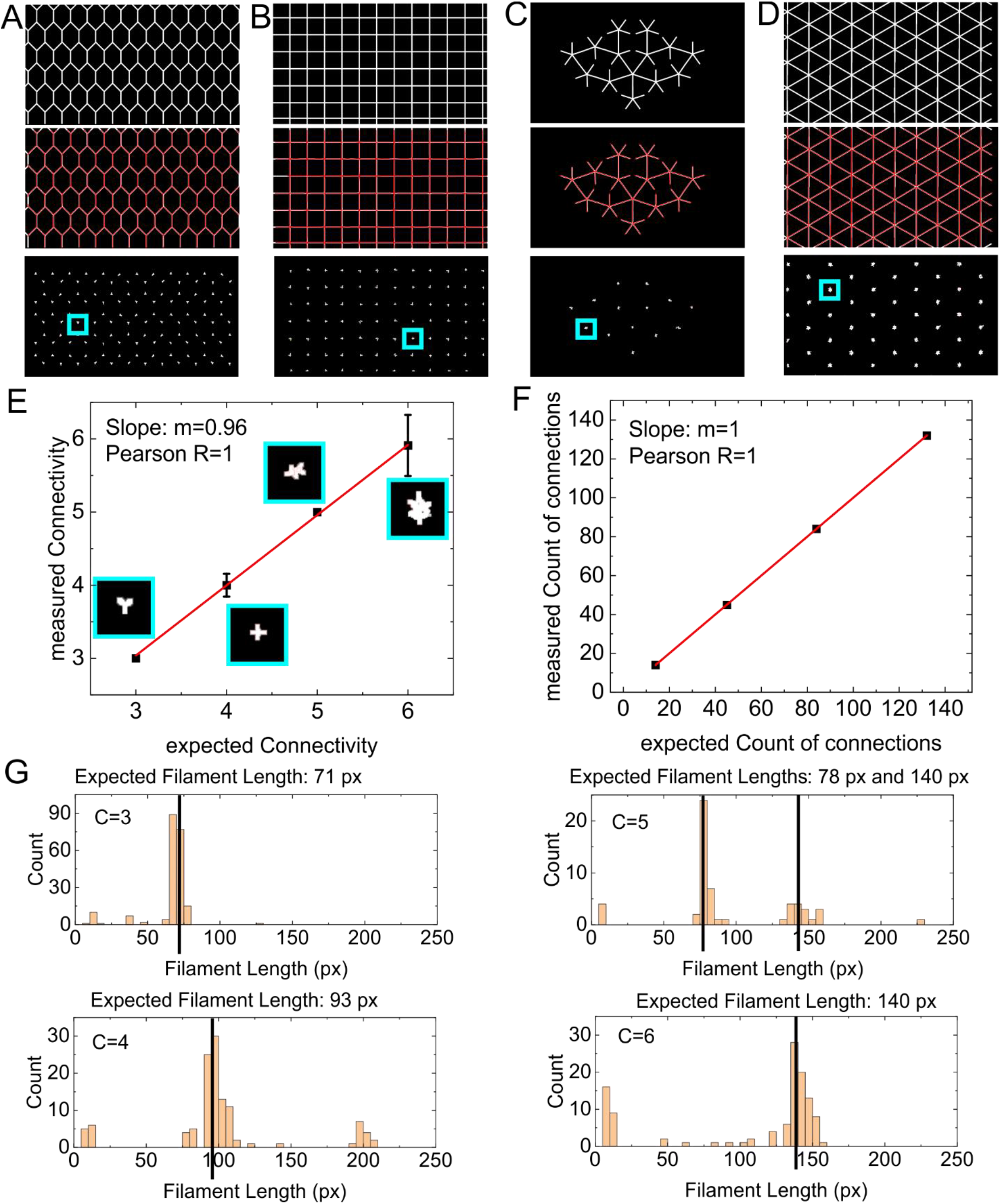
Connectivity and Filament Length length tests. (**A-D**) Exemplary images with discretely increasing Connectivities as three (A), four (B), five (C), and six (D), as are traced in red (second row), and with branches/junctions extracted (third row). Junction Connectivity was set to 6 (i.e., 6 nodes connected to identify a single fiber belonging to one junction). Line thickness: 9 px. (**E, F**) Linear approximations of expected *versus* measured Connectivity (insets show corresponding third rows of (A-D)) (E) and Counts of branches/connections (F). Red lines show the linear approximation. (**G**) Filament Length histograms of images (A-D) corresponding to the four Connectivities from three to six. The break-off angle was set to ~17° (0.3 rad). Black vertical lines represent the expected Filament Length.

### 3.4. Filament Length

In addition to the Mesh Hole Size and Connectivity, the Filament Length is a crucial parameter in the characterization of networks. To include this parameter in the FiNTA, a user-defined break-off angle was implemented to detect and define a filament as a single filament (see Fig. 1C).

To determine the accuracy of this Filament Length tracing approach, the artificially created test images for the Connectivity were modified to show a homogenous distribution of Filament Lengths by removing the filaments at the edges of the images. This resulted in images with very similar Filament Lengths, as shown in Figure 5G for a break-off angle of ~17° (0.3 rad), where all of the Filament Lengths were within an accurate range of the expected lengths. However, the FiNTA also identified artificial filaments that were significantly shorter than those expected, and were localized to the branching/connection zones. Such very short filaments can be eliminated easily using a threshold value. We also performed a more detailed test of the Filament Lengths and their dependence on the break-off angle, as presented in Figure S1.

### 3.5. Other extracted parameters

As the Mesh Hole Size, Connectivity and Filament Lengths should be the most challenging parameters to implement, we mainly focused on testing these parameters in detail. Additional parameters that can be calculated based on the Mesh Hole Size and from the tracing *per se* were also tested. The filament density can be calculated from the length of the total traced network divided by the image size. The Junction Distance was calculated forwards from the known junction positions. The Connectivities from the FiNTA were very accurate for both the quality and the counts, and thus the reliability of the analysis of Junction Distance was also accurate, as for the Connectivity. The Circularity is the fractal dimension of the Mesh Holes, and this was defined as proportional to the Mesh Hole Size divided by the perimeter squared. The reliability of the analysis of Circularity is therefore the same as that for the Mesh Hole Size, which was shown to be very accurate in section 3.1.

### 3.6. Tracing accuracy

The accuracy of the tracing determines the quality of all of the parameters that are extracted from any tracing software. To determine the tracing accuracy with the FiNTA, we analyzed electron microscopy images of actin filaments and a fluorescence confocal microscopy image of microtubules (Fig. 6). For the quality of the analysis, the filaments traced with the FiNTA were then also traced manually in the same images. This thus determined the relative levels of false-negative and false-positive signals obtained using the FiNTA, as the relative proportions of nontraced filaments and nonexisting filaments traced, respectively.

**FIGURE 6.**
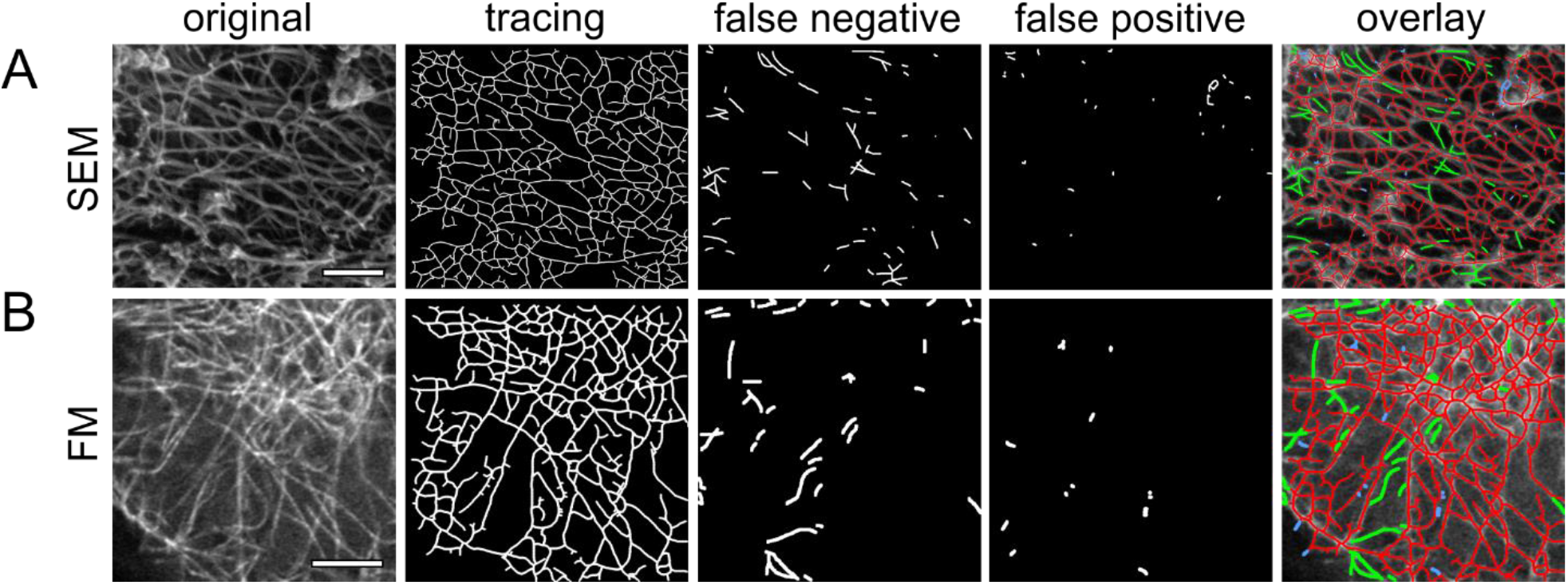
Tracing error tests. Representative images of actin filaments obtained using scanning electron microscopy (**A**; SEM) and of microtubules obtained using confocal fluorescence microscopy (**B**; FM), as traced with the FiNTA and compared to hand tracing. Nontraced fibers (false negative) and nonexisting filaments (false positive) that were traced are shown (2× fiber thickness of original tracing). Overlay of the original tracing (red), false negatives (green), and false positives (blue) shown in the last column. Scale bars: 200 nm (A); 5 μm (B).

For determination of the levels of false-negative signals, the relative numbers of nontraced filaments were calculated by dividing these by the total network length traced by hand including the nontraced filaments. For determination of the levels of false-positive signals, the proportion of false-positive filaments were defined according to the total network length traced by hand. On this basis, the FiNTA tracing for the electron microscopy images showed 8.55% false-negative filaments and 1.54% false-positive filaments. For the fluorescence image FiNTA tracing, there were 9.72% false-negative filaments, and 1.42% false-positive filaments. Therefore, the false-positive rates can be considered as negligible, and the false-negative rates are <10%, which can be considered an acceptable value. The false-negative filaments arise because the FiNTA does not trace up to the edge of the image, to avoid any distortion of the Mesh Hole Size. The rates of false-negative tracing can therefore be reduced by cropping the images. In the central parts of the images, almost all of the filaments that are not traced (i.e., the false negatives) arise where the angle between two filaments is small.

### 3.7 Network separation

It is challenging to draw conclusions about spatial distributions in three dimensions from 2D images of 3D networks, both manually and automatically. For example, it is usually not possible to distinguish between a real junction and an overlay of two filaments. However, the FiNTA can reduce these events to a minimum. Here, this is based on the effect whereby the filaments closer to the camera (i.e., at a higher level within the network) are brighter than the filaments that are further away from the camera (i.e., at a lower level within the network). This is true for images from both electron microscopy and fluorescence microscopy. For this, the option was implemented to preserve traced nodes within a specified grayscale interval only, which can be chosen by the user. This approach can also be used to avoid over-exposed and under-exposed regions of a network.

To assess the quality of this approach, three different images were analyzed at high and low intensity levels (Fig. 7). For electron microscopy images of actin networks, the high intensity areas, referred to as ruffles, provide no information about the network due to their overexposure. It might therefore be of interest to exclude these high intensity regions from the tracing, which is what the FiNTA does (Fig. 7A). To determine the quality of the FiNTA for other types of networks, an electron microscopy image of the hexagonal structure of a diatom alga was analyzed. The thresholding of the intensity values allowed the FiNTA to discern the hexagonal structures from the underlying network (Fig. 7B). Furthermore, in fluorescence images of actin, the FiNTA can differentiate between high intensities (more fluorescein proteins) and low intensities (less fluorescein proteins) (Fig. 7C). By using intensity thresholding, the FiNTA can split a grayscale image into all of the gray levels. This allows the user to focus on the subcellular regions of a network, which can be useful if the user is interested in subregions, or parts of the intensity distribution within an image. Therefore, such intensity thresholding is a powerful method to separate networks for their detailed investigation.

**FIGURE 7.**
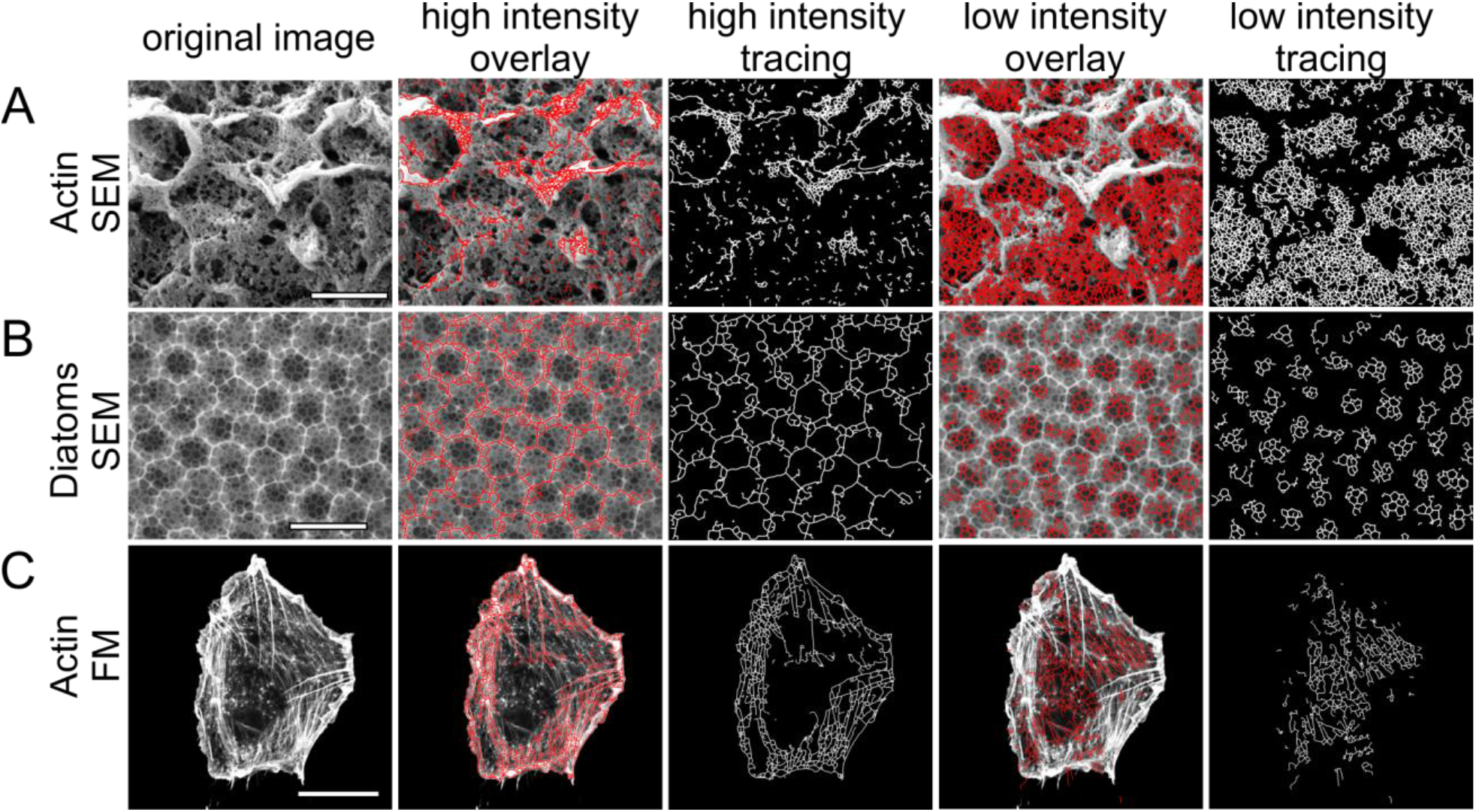
Network separation by intensity thresholding. Representative images for scanning electron microscopy (SEM) of actin (**A**) and a diatom (**B**) and fluorescent microscopy (FM) of actin (**C**), used to separate the networks by intensity. (**A, B**) As used to eliminate unwanted areas (A, actin: high intensity) and analyze the remaining network (low intensity), or to separate a superior (B, diatom, high intensity) and an inferior (low intensity) network. (**C**) As used to separate between high and low intensity signals that represent high and low amounts of fluorescent actin molecules. Scale bars: 1 μm (A), 4 μm (B), 20 μm (C).

### 3.8. Tracing of cytoskeletal networks imaged by electron and fluorescence microscopy

To demonstrate the wide range of FiNTA applications, images of different cytoskeletal filaments were traced that were obtained by electron microscopy (Fig. 2) and using different variants of fluorescence microscopy (Fig. 8).

**FIGURE 8.**
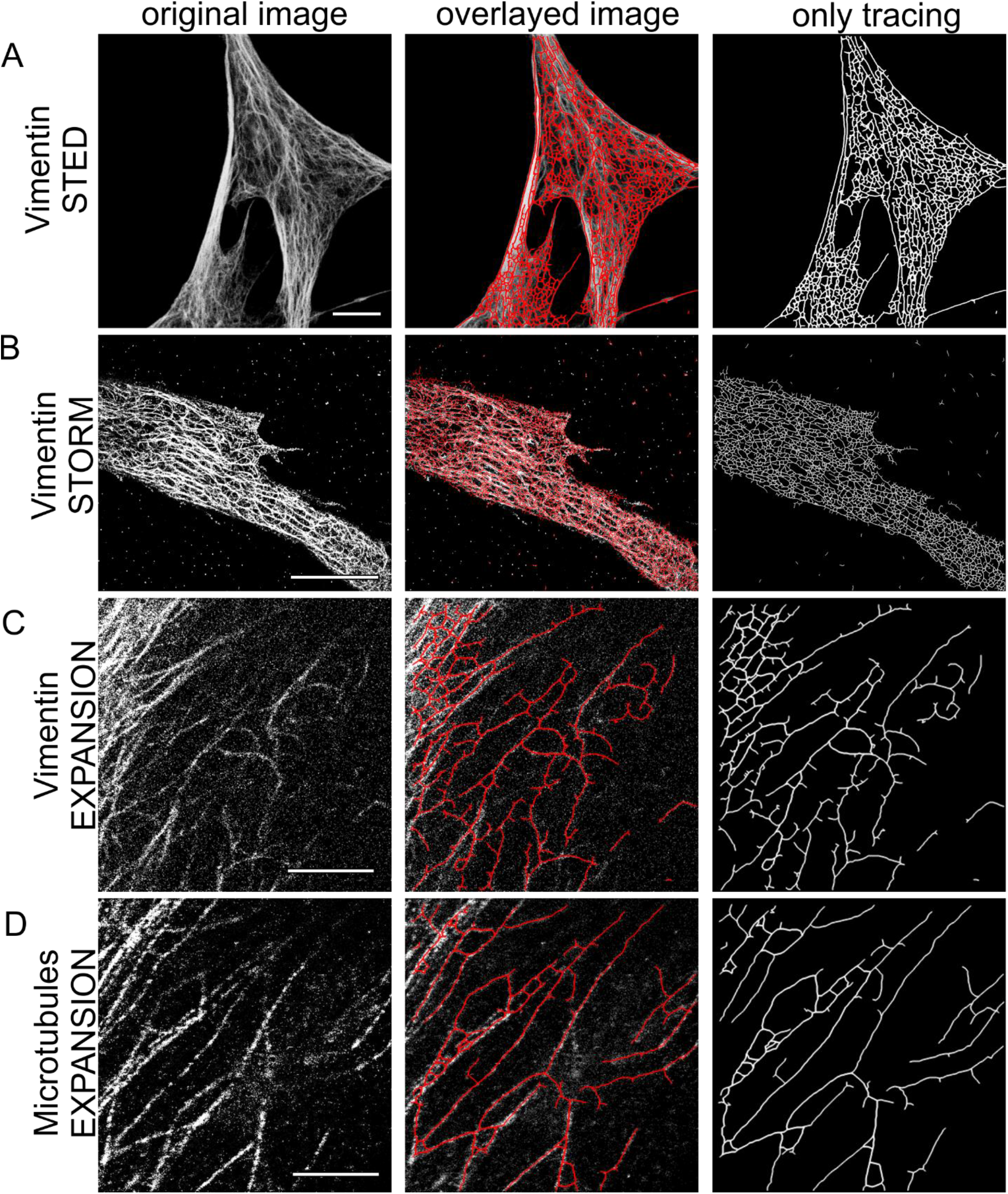
Representative images and exemplary traced fluorescence super-resolution networks. Vimentin was imaged and traced (as indicated) using stimulated emission depletion (STED) microscopy (**A**), 3D stochastic optical reconstruction microscopy (STORM) (**B**) and Expansion microscopy (**C**). (**D**) Identical image section of cell vimentin in (C) is shown for microtubules in (D). Overlayed image: overlay of original and traced images. Scale bars: 10 μm (**A, B**) (expansion factor, 4.5); 2.38 μm (**C, D**).

We showed in section 3.7. that the FiNTA can be used to analyze actin microfilament cytoskeletal networks imaged by confocal microscopy (Fig. 7C). However, the resolution of confocal microscopy is often not sufficient to determine the fine structure of cytoskeletal networks, as more recently, super-resolution techniques are being increasingly used. To provide a useful tool for such studies, the reliability of the FiNTA was analyzed using images from super-resolution microscopy. Images of the intermediate filament protein vimentin were analyzed for various cell types that were provided by three distinct super-resolution microscopy techniques that are based on different approaches and principles: Stimulated Emission Depletion (STED) microscopy; Stochastic Optical Reconstruction Microscopy (STORM); and Expansion microscopy. STED microscopy is a type of confocal microscopy where a homogenous signal along filaments is recorded. In contrast, STORM and Expansion microscopy are techniques that create a pointillism-like signal along filaments. In STORM microscopy, this is achieved by the imaging technique itself, whereas in Expansion microscopy, the structure under investigation is physically ruptured at the nanoscale level. These vimentin network images using each of these super-resolution techniques were indeed accurately traced by the FiNTA (Fig. 8A-C). This included the pointillism-like signals of STORM and Expansion microscopy, as shown in the detailed pointillism test in Figure S2, and more indirectly in Figure 4. To further confirm this, Figure 8D shows the FiNTA tracing of a microtubule meshwork obtained using Expansion microscopy. Consequently, we can conclude that the FiNTA can also correctly trace networks in images using super-resolution fluorescence imaging techniques.

As indicated in section 2.2., the FiNTA can trace actin microfilament networks obtained using electron microscopy. To determine whether the FiNTA can also be used for electron microcopy images of other structures, we additionally investigated mushroom braid, eggshell, foam, and diatoms. With the exception of the foam, which showed high heterogeneity for the filament thickness and brightness, the FiNTA provided accurate tracing of these networks (Fig. S4).

To further validate the FiNTA, other images were analyzed, including a picture of sticks of spaghetti obtained using a reflex camera, and different painted, biological and written images, which again resulted in consistently accurate tracing (Fig. S5).

## 4. Discussion

In this study, we present the powerful, rapid, user-friendly, open-source vectorial FiNTA for grayscale images of different filament types and across a large variety of scales. This is shown by our analysis of the intracellular nanoscale actin microfilaments and vimentin cytoskeleton, and of other network forms using various imaging techniques.

A lot of the available software and plugins for analysis of networks require the raw data images in grayscale to be converted into binary images prior to analysis, such as the ImageJ plugins DiameterJ and NeuronJ. The quality of the images will thereby greatly impact on the recognition of filaments by the software. With the FiNTA, grayscale images can be analyzed, which thus overcomes this problem.

The FiNTA allows analysis of the common network parameters, such as Mesh Hole Size, Circularity, Junction Distance, Filament Density and others. Moreover, it provides a high level of quantification of challenging parameters, such as Connectivity and Filament Length. The FiNTA can also be used to separate sub-networks in single images, through intensity thresholding. It has a high tolerance in terms of filament density and noise. Moreover, the FiNTA delivers good results regardless of the imaging technique used (e.g., fluorescence microscopy, scanning electron microscopy, photographs), and of the substance investigated.

One limitation of the FiNTA relates to networks with small angles between the filaments. The FiNTA was also implemented for homogenous filament thickness. Although the FiNTA can be used where there are small variations in filament thickness, significant heterogeneity in filament thickness across a network can lead to nonideal tracing. Finally, heterogeneous intensity distributions along filaments can result in increased false tracing by the FiNTA. In such cases, segmentation-based algorithms might provide more accurate tracing.

In summary, the FiNTA is an algorithm that can provide great benefits across a wide variety of scientific fields that involve analysis of images of networks.

## Acknowledgement

The authors would like to thank the Leibniz Institute for New Materials (INM), Saarland University, and the DFG (CRC 1027 (A10)) for financial support. The authors would also like to thank the Weston Park Cancer Centre (University of Sheffield) the Fundação para a Ciência e a Tecnologia (FCT), the Portuguese Government (PEst-OE/QUI/UI0674/2013) and the Agência Regional para o Desenvolvimento da Investigaçaõ Tecnologia e Inovação (ARDITI), M1420-01-0145-FEDER-000005-Centro de Química da Madeira—CQM (Madeira 14-20). We thank Gaudenz Danuser (Texas, USA) for the hTERT-RPE1 cells stably expressing mEmerald-vimentin and mTagRFPt-α-tubulin, Matthieu Piel (Paris, France) for the hTERT-RPE1 cells stably expressing mCherry LifeAct, and William C. Hahn (Harvard Medical School, USA) for the BJ cell line. We thank Ana Lopez and Christa Walther, University of Sheffield, UK, for excellent technical support. We thank Guillaume Charras (London Centre of Nanotechnology, UK) for excellent ideas about the parameters extracted by the FiNTA.

## Author contributions

DF, MS and ET designed the study

MS developed the algorithm

DF and MS tested the algorithm

DF and MK prepared the samples and provided the electron microscopy images, which were obtained at the Leibniz Institute for New Materials (INM), Saarbruecken, Germany.

AG prepared samples and provided the STORM images, which were obtained at The Wolfson Light Microscopy Facility & University of Sheffield, UK.

DT prepared samples and provided fluorescent STED images of vimentin, which were obtained at the Leibniz Institute for New Materials (INM), Saarbruecken, Germany.

LK prepared samples and provided fluorescent images of microtubules, which were obtained at the Leibniz Institute for New Materials (INM), Saarbruecken, Germany.

DF prepared samples and provided Expansion microscopy images of microtubules and vimentin, which were obtained at the Leibniz Institute for New Materials (INM), Saarbruecken, Germany.

DF, AG and FL wrote and revised the manuscript.

## Supporting information

### 1. Algorithm method

#### 1.1. Image analysis using the filament network-tracing algorithm

The aim of the filament network-tracing algorithm (FiNTA) is to trace filaments on grayscale images with increased precision and detail compared to classical segmentation-based algorithms. This was achieved by populating the images with variable numbers of starting nodes on the filaments. A circle was taken around each node. The FiNTA defines where the filaments are localized on the circle, using the second-order derivative of the Hessian matrix. At this location, a second node connected to the first one is set. This procedure is performed for every node, until the whole network is traced. In this way, each node can be connected to an arbitrary number of other nodes, which allows the branching of the traced network and the formation of closed loops.

#### 1.2. Details of the tracing algorithm

Before the image convolution by the FiNTA, the image is converted into an 8-bit image. To compute the smoothed second-order derivatives of the image brightness, the tracing algorithm works on the basis of the Hessian matrix of the Gaussian transform. The convolution with the three second-order derivatives of the Gaussian transform yields the Hessian matrix for each pixel. Using the Hessian matrix, the directional second-order derivative for any pixel and any direction can then be computed. Generally, negative second-order derivatives with large absolute values indicate the presence of a filament. This filament travels perpendicular to the direction of the directional second-order derivative. An automatic function sets several starting nodes. From these nodes, the FiNTA traces the whole network without multiple tracing of the filaments. The FiNTA is available at: https://github.com/SRaent/Actin.

#### 1.3. Input parameters

One key feature of the FiNTA is that it is highly adaptable to a wide range and scale of images of networks. The eight input parameters create the flexibility that allows the user to adjust the FiNTA to a specific network of interest. Another advantage is that there is no need to define and potentially optimize any of the parameters choices, because the parameters are defined as in Table S1, with their suggested starting values that generally lead to correct tracing, for almost all cases investigated. Finally, the consequence of over-estimation and under-estimation of the individual parameters are also described in Table S1.

### 2. Further algorithm testing

#### 2.1. Details for Filament Length and Connectivity

To quantify the accuracy of the Filament Length tracing by the FiNTA, the artificially created Connectivity images were modified such that no filament reached the edge of the image, to create a homogenous Filament Length distribution. The original images, traced images, and resulting Filament Lengths are illustrated in Figure S1A-D. Each of the different colors corresponds to a slightly different Filament Length, which depends on the total Filament Length distribution. The Filament Length calculated by the FiNTA strongly depend on the break-off angle set by the user. Therefore, the length histograms are shown for the four investigated images from 6° to 115° (0.1-2.0 rad) in Figure S1E-H. Independent of the angles within the investigated networks, the optimal break-off angles were between 17° and 26° (0.30-0.45 rad). This universal range of break-off angles that lead to the correct Filament Lengths can be reasoned on the basis that the angle between each of two connected nodes defines the break-off angle. Consequently, the angle between the filaments is independent of the break-off angle for the calculation of the Filament Length. The drawback of this method is that it increases the short false filaments, which occur at the junctions. In addition, the junctions can be ignored as intended, which leads to Filament Lengths that are multiples of the histogram peaks. Consequently, we suggest that the resulting histograms are always investigated carefully and the Filament Length optimized by thresholding. The analogous argumentation is valid for the Filament Count, as demonstrated in Figure S1I-L.

Despite the explained weaknesses of the Filament Length calculations, the FiNTA can measure the real Filament Length independent of the Junction Distance. We conclude that the Filament Length calculation by the FiNTA has to be further investigated by hand for more accurate results, although the histogram analysis provides correct results.

Junctions close to each other frequently result in incorrect Connectivities. Therefore, we implemented summarization of junctions to one junction. For this, the user needs to specify a unification distance. The FiNTA unites all of the junctions that can be connected by the number of steps specified by the unification distance. These is interpreted as one junction. One step is considered to be the distance between two nodes that are connected to each other.

#### 2.3. Pointillism tests

Many algorithms are dedicated to the analysis of images using a specific technique or even from a specific network. In contrast, the FiNTA can be used to analyze networks imaged with any of the existing imaging techniques, as long as the filament thickness and brightness are relatively homogenous (see filament thickness in section 2.4. below) and the image type is grayscale.

Some imaging techniques result in pointillism-like signals, such as STORM. Depending on the sampling and post-processing, the distribution of signal points can vary. We tested the tolerance of the FiNTA to pointillism by creating images with increased black pixels on white filaments, analogous to the noise tests in Figure 4. Therefore, we increased the amount of black pixels on the white filaments from 0% to 100% (Fig. S2A). In addition, we varied the line thickness (see Fig. S4), and used the Mesh Hole Size and Count as quantification parameters. By calculating the percentage errors between the measured and expected Mesh Hole Size Sizes and Counts, we show that the FiNTA traces pointillism filaments very accurately (Fig. S2B, C). The larger the filament thickness, the more pointillism is accepted by the FiNTA. We conclude that the FiNTA can be used for any imaging technique that creates pointillism-like structures of networks.

#### 2.4. Filament thickness

Initially, the FiNTA was implemented to quantify filament networks with uniform filament thicknesses. Nonetheless, the FiNTA has a tolerance regarding heterogeneous filament thicknesses. For quantification of this tolerance, an image was artificially created with 13 lines of different filament thicknesses, from 1 px to 13 px, in 1 px steps. Each line was connected to the following thicker line once. The connection line had the thickness of the thicker line (see Fig. S3A). In total, 29 different filament thicknesses were assigned to the FiNTA, from 2 px to 30 px, in 1 px steps. Figure S3B-E provides examples for assigned filament thicknesses of 2 px, 10 px, 20 px, and 30 px. Three parameters were tested here: recognition of filament thickness; accuracy of the traced filament thickness; and accuracy of the traced connection thickness.

The filaments recognized by the FiNTA are defined as the filaments that are traced at least for 90% of their full length. The tracing line can be anywhere on the filament (including at the edges), while double tracing of one filament is prevented. Here, the FiNTA can handle roughly ±100% of the assigned filament thicknesses (Fig. S3F).

The accurately traced filament thickness is defined as the recognized filament thickness, under the requirement that the tracing line is not on the edge of a filament (except when close to the connection). The FiNTA leads to correct tracing of up to 100% thicker filaments than the assigned filament thickness, and about 70% to 80% thinner filaments than the assigned filament thickness (Fig. S3G).

The accurately traced connection thickness is defined as the filament thickness of a connection filament for which neither multiple filament tracing nor tracing of the background occurs. The FiNTA leads to correct tracing of roughly 20% thicker connection filaments than the assigned filament thickness above a 5-px filament thickness (Fig. S3H). Below the 5-px filament thickness, more or less none of the filaments thicker than the assigned filament thickness were correctly traced. In contrast, the FiNTA leads to correct tracing of filaments that are between 60% and 86% thinner than the assigned filament thickness (the only exclusion was an assigned filament thickness of 2 px, because the smallest filament was 1 px thick). Consequently, although depending on the heterogeneity of the filament thickness of interest, if all of the filaments are to be traced, we would recommend to tend to over-estimation of the filament thickness than to under-estimation. In contrast, this effect can be very useful to discern between filaments of different thicknesses, as long as the difference is in the range of the filament thickness itself.

### 3. Biological material and methods

#### 3.1. Cell culture

Immortalized retinal pigmented epithelium (hTERT-RPE1) cells were grown in Dulbecco’s modified Eagle’s medium (DMEM)/F12 medium supplemented with 10% fetal bovine serum (Thermo Fisher, MA, USA), 1% glutamax, and 1% penicillin/streptomycin under 5% CO2 at 37 °C, in cell culture flasks (Cellstar, Greiner Bio-One, Austria).

For fluorescent imaging, hTERT-RPE1 cells stably expressing mEmerald–vimentin and mTagRFPt–α-tubulin (a kind gift from Gaudenz Danuser, Dallas, TX, USA) or mCherry LifeAct (a kind gift from Matthieu Piel, Paris, France) were placed in glass-bottomed dishes (Fluorodish; World Precision Instruments, Germany) and left to adhere overnight. Live cells were imaged at 37 °C and 5% CO2 using laser scanning confocal microscopy (LSM 980; Zeiss, Germany).

Bjhtert SV40T V12 H-Ras^1^ cells were cultivated at 37 °C in 5% CO2 in DMEM (Sigma-Aldrich, Germany) supplemented with 10% fetal bovine serum (VWR, Germany) and 1% antibiotic/antimycotic solution (Invitrogen, Germany). The cells were harvested at approximately 30% confluence.

Baby hamster kidney (BHK-21) cells were cultured in DMEM/F12 supplemented with 5% fetal bovine serum and 1% penicillin/streptomycin under 5% CO2 at 37 °C.

### 4. Fluorescence protocols and imaging

#### 4.1. Fluorescence microscopy

For fluorescent imaging, RPE1 cells that stably expressed mCherry LifeAct (a kind gift from Matthieu Piel, Paris, France) were fixed in 0.5% paraformaldehyde and mounted with Fluomount-G mounting medium, with DAPI (FisherScientific, Germany). If not stated otherwise, the imaging was performed using laser scanning confocal microscopy (LSM 980; Zeiss, Germany).

#### 4.2. Stochastic optical reconstruction microscopy

Bjhtert SV40T H-RasV12 cells were seeded onto glass coverslips that were pre-cleaned, as described previously^2^. After 48 h, the cells were washed in phosphate-buffered saline (PBS) at 37 °C, and the autofluorescence and paraformaldehyde were quenched. The cells fixed in PBS with 3.7% paraformaldehyde and 0.2% Triton X-100 for 15 min at 37 °C, blocked, and immunostained with a mouse anti-vimentin antibody (V9; 1:50), followed by anti-mouse IgG Alexa 647 (1:400), as described previously^2^. GLOX buffer was used, where 1 mL contained: 0.1 g glucose (in 10 μL), 100 μL glucose oxidase (at 0.005 g/mL), 100 μL cysteamine/MEA (at 1 M cysteamine/MEA), in 790 μL Tris/NaCl buffer (at 50 mM Tris, with 10 mM NaCl). Images were taken with a 20-ms exposure time for the N-STORM (Nikon), for a resolution of ~50 nm.

#### 4.3. Stimulated emission depletion microscopy

BHK-21 cells were plated on coverslips coated with poly-l-lysine. The cells were fixed with ice-cold acetone for 10 min at −21 °C. The samples were washed with PBS, blocked using 5% goat serum in PBS for 1 h, and incubated with the anti-vimentin antibody (V9; 1:200) for 1 h at room temperature. After a further wash with PBS, the samples were incubated with Abberior Star Red (1:500) for 1 h. The coverslips were washed with PBS and mounted using Mowiol, for 12 h at room temperature. Image acquisition with STED microscopy was then performed (SP5-STED; Leica, Germany).

#### 4.4. Expansion microscopy

The preparation for Expansion microscopy was performed following the protocol for cell culture of Tillberg and co-workers^3^. In brief, hTERT-RPE1 cells stably expressing mEmerald–vimentin and mTagRFPt–α-tubulin were fixed in 0.5% paraformaldehyde. Anchoring with Acyloyl-X was followed by *in-situ* polymerization with a hydrogel. The addition of water after the digestion (including with proteinase K) physically increased the sample size by a factor of 4.5, in all spatial directions.

### 5. Scanning electron microscopy preparation and imaging

All of the scanning electron microscopy images were obtained using an environmental scanning electron microscope (Quanta 400; FEI, USA).

#### 5.1. Actin cortex

The preparation of the actin cortex of hTERT-RPE1 cells for the electron microscopy was similar to the protocols of Svitkina, Chugh, and others^4, 5^. In brief, the cell membrane was disrupted with Triton X-100, and the cells were lightly fixed, at the same time. After stronger fixing with glutaraldehyde and other fixation agents, the cells were dehydrated using a series of increasing ethanol concentrations. Hexamethyldisilazane was used as the final drying procedure before the samples were sputtered with 6 nm to 7 nm platinum. Images were taken at 5 kV under high vacuum conditions, using secondary electrons, with an Everhart–Thornley detector.

#### 5.2. Mushroom braid

Wet mushroom braid was dried under ambient conditions and visualized at 10-kV accelerating voltage under low vacuum conditions (water vapor, 100 Pa) using secondary electrons, with a large field detector.

#### 5.3. Eggshell

A piece of an egg shell was placed on double-sided carbon tape with the outer surface on top. Secondary electron imaging using a large field detector was performed at 10-kV accelerating voltage under low vacuum conditions (water vapor, 100 Pa).

#### 5.4. Foam

A small piece of foam packaging was cut using a blade and fixed using double-sided carbon tape. The surface analysis was carried out at 5-kV accelerating voltage, under low vacuum conditions (water vapor, 100 Pa), using secondary electrons, with a large field detector.

#### 5.5. Diatoms

Diatoms were placed on double-sided carbon tape and imaged at 10-kV accelerating voltage under high vacuum conditions using secondary electrons, with an Everhart–Thornley detector.

---

**TABLE S1.** Overview of the parameters assigned to the filament network-tracing algorithm (FiNTA).

| Parameter | Definition | Starting value |  |  |
| --- | --- | --- | --- | --- |
|  |  | Suggested | Too low | Too high |
| ***$\sigma_{conv}$*** | Image smoothing and Hessian matrix is computed | Half the fiber diameter, in pixels | Multiple tracing of single fibers | Not all fibers traced |
| ***$\sigma_{smooth}$*** | Smoothing of angle-dependent curvature of the fibers | 0.5 radians (0.35 < $\sigma_{smooth}$ < 0.8) | Too many nodes are generated | Not enough branches, with small angles between fibers |
| ***$r_{min}$*** | Minimal distance between nodes | 0 px (< half fiber diameter) | -- | No tracing |
| ***$r_{max}$*** | Maximal distance between nodes | Fiber diameter in pixels | Many branches are not traced | Influence of surrounding fibers on tracing, and increased computation time |
| ***$r_{step}$*** | Distance between generated and new nodes | Half fiber diameter < $r_{step}$ < fiber diameter, in pixels | Single fibers are traced more than ones, and increased computation time | Branches are not found, thus decreased tracing smoothness |
| ***$steps$*** | Number of angles for which new nodes can be generated | 100 < steps < 360 | Minor changes in tracing quality | Minor changes in tracing quality, and linear increase in computation time |
| ***$thresh$*** | Value above which the local maxima in the smoothed angle depending curvature are used to generate nodes | 0.5 | A lot of nodes in noisy areas | No nodes generated |
| ***$min\_loop\_length$*** | Minimal length of loops, expressed as number of nodes belonging to a loop | 5 < $min\_loop\_length$ < 10 | Unwanted connections | Increasing number of almost closed loops |

**FIGURE S1.**
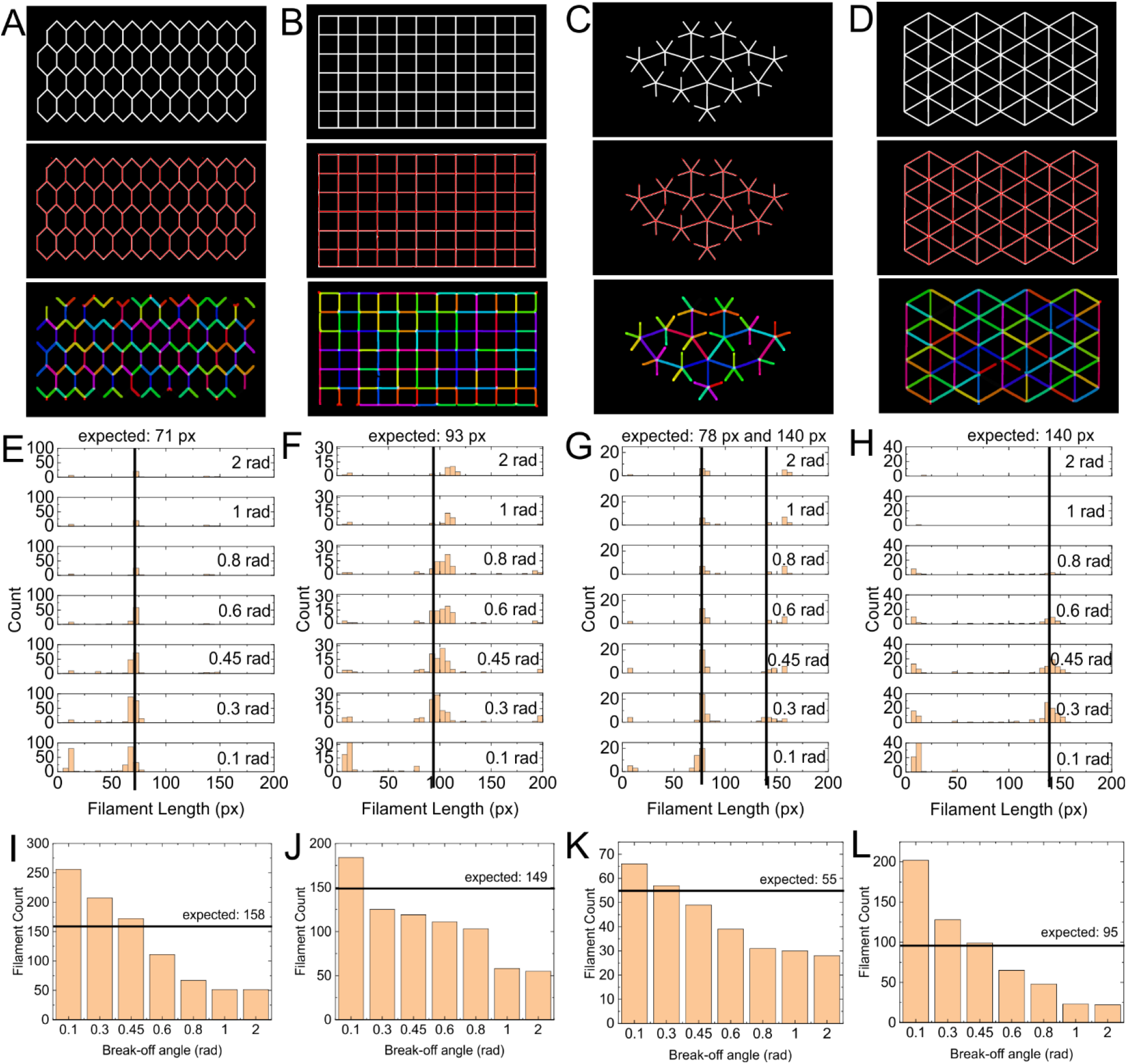
Filament Length tests. (**A-D**) The exemplary images used for the Connectivity tests (Fig. 4) were also used for these length tests, because of the high length homogeneity within each image. The traced images (second row) and color-coded lengths (third row) are also shown. The color coding was applied to better differentiate between the Filament Lengths. (**E-L**) Quantification of filament lengths (E-H) and filament counts (I-L) for each of the images in (A-D), respectively. Black lines, expected Filament Lengths (E-H) and Filament Counts (I-L).

**FIGURE S2.**
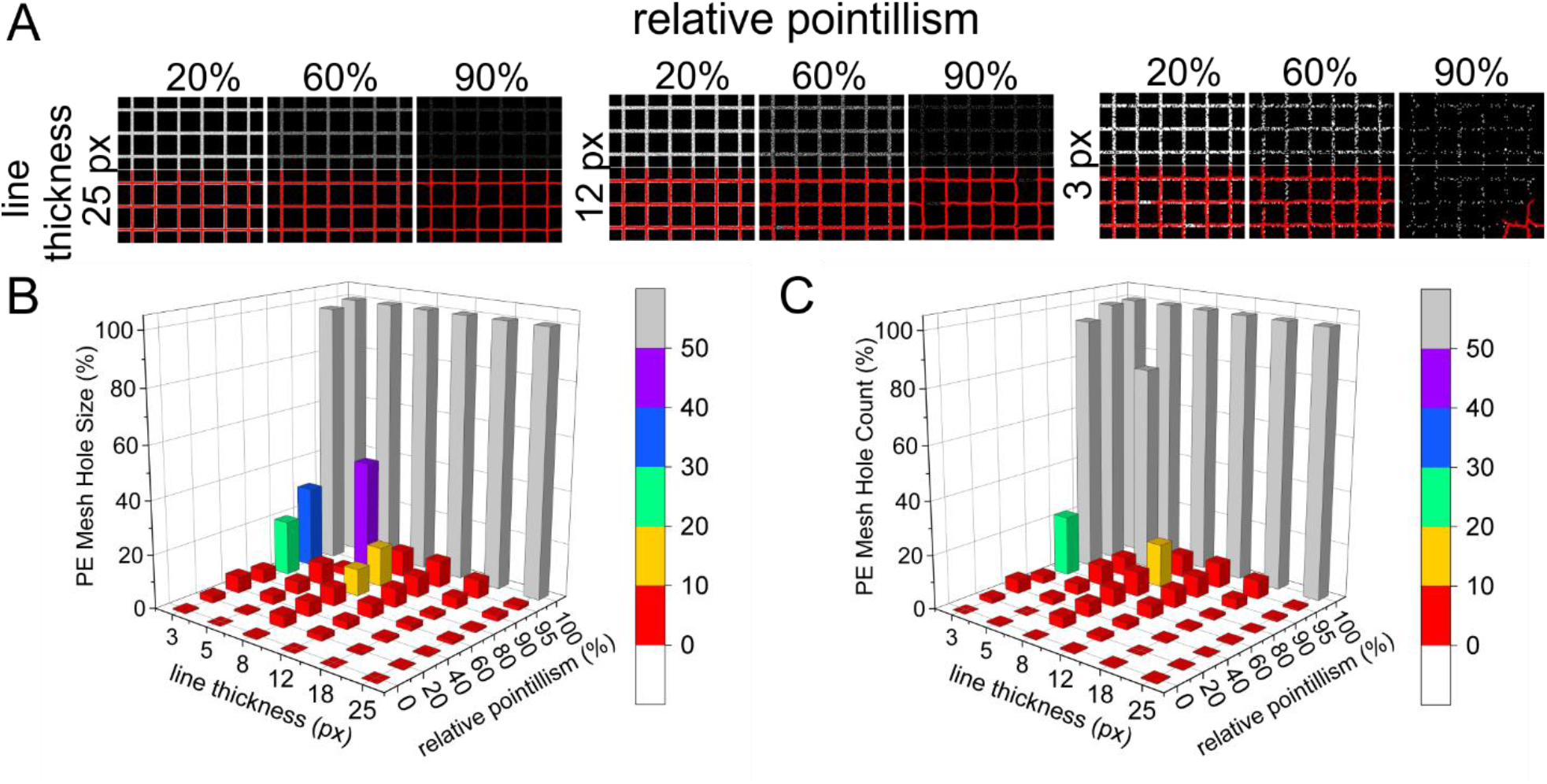
Mesh Hole Size determination by pointillism and line thickness. (**A**) Exemplary images of the traced Mesh Holes for the varied pointillism (amount of black pixels on white lines) and line thickness. (**B, C**) Percentage errors (PE) of the expected Mesh Hole Size (B) and Counts (C) *versus* the measured values, according to line thickness and signal-to-noise ratio.

**FIGURE S3.**
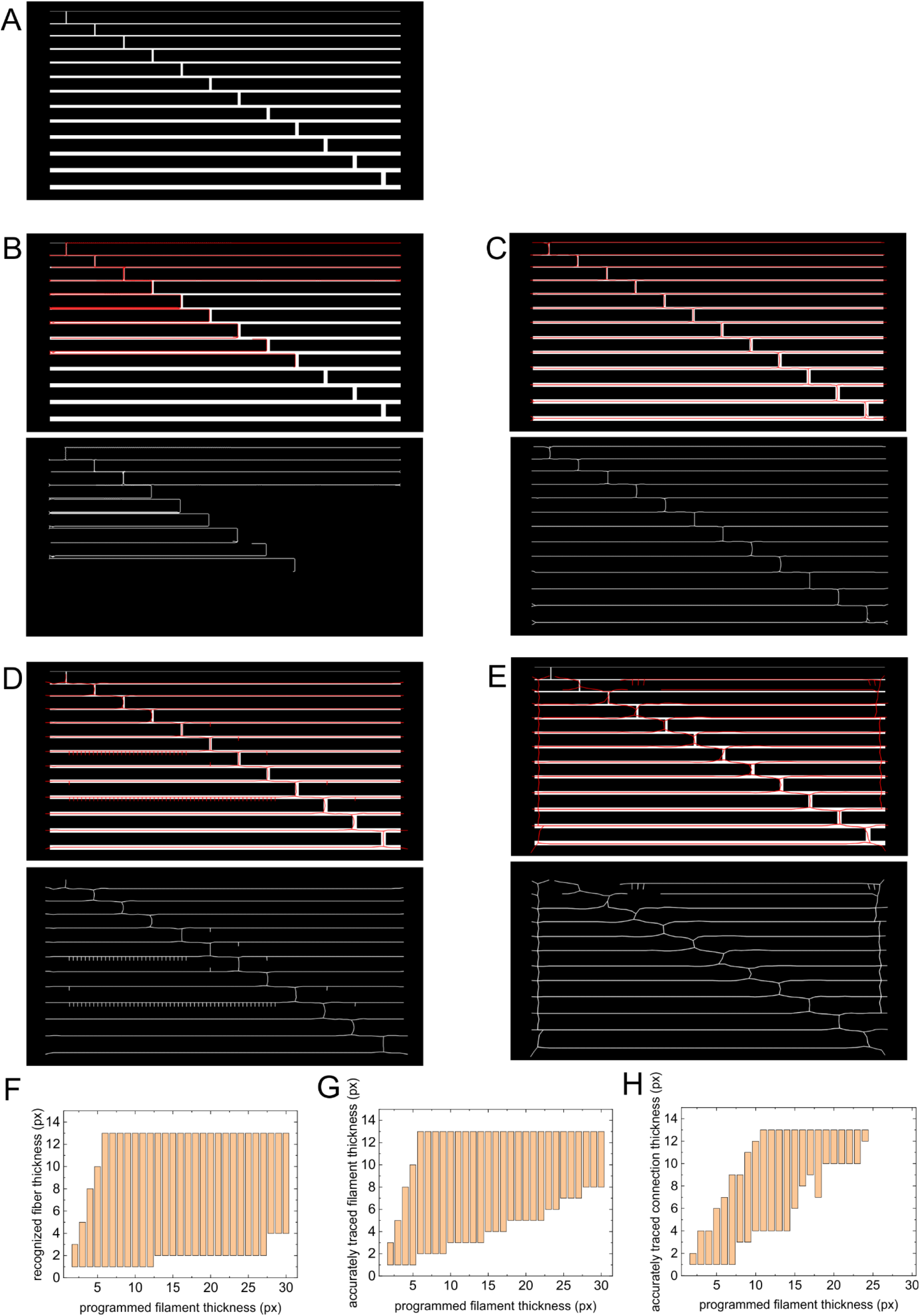
Filament thickness tests. (**A-E**) Exemplary images with lines ranging from 1 px to 13 px in 1-px steps (A), and their tracing (B-E). Each line is connected to the next thicker line by a connection with the thickness of the thicker line. (**F-H**) Quantification of corresponding recognized fiber thickness (F) and the accurately traced fiber thicknesses (G) and connections (H).

**FIGURE S4.**
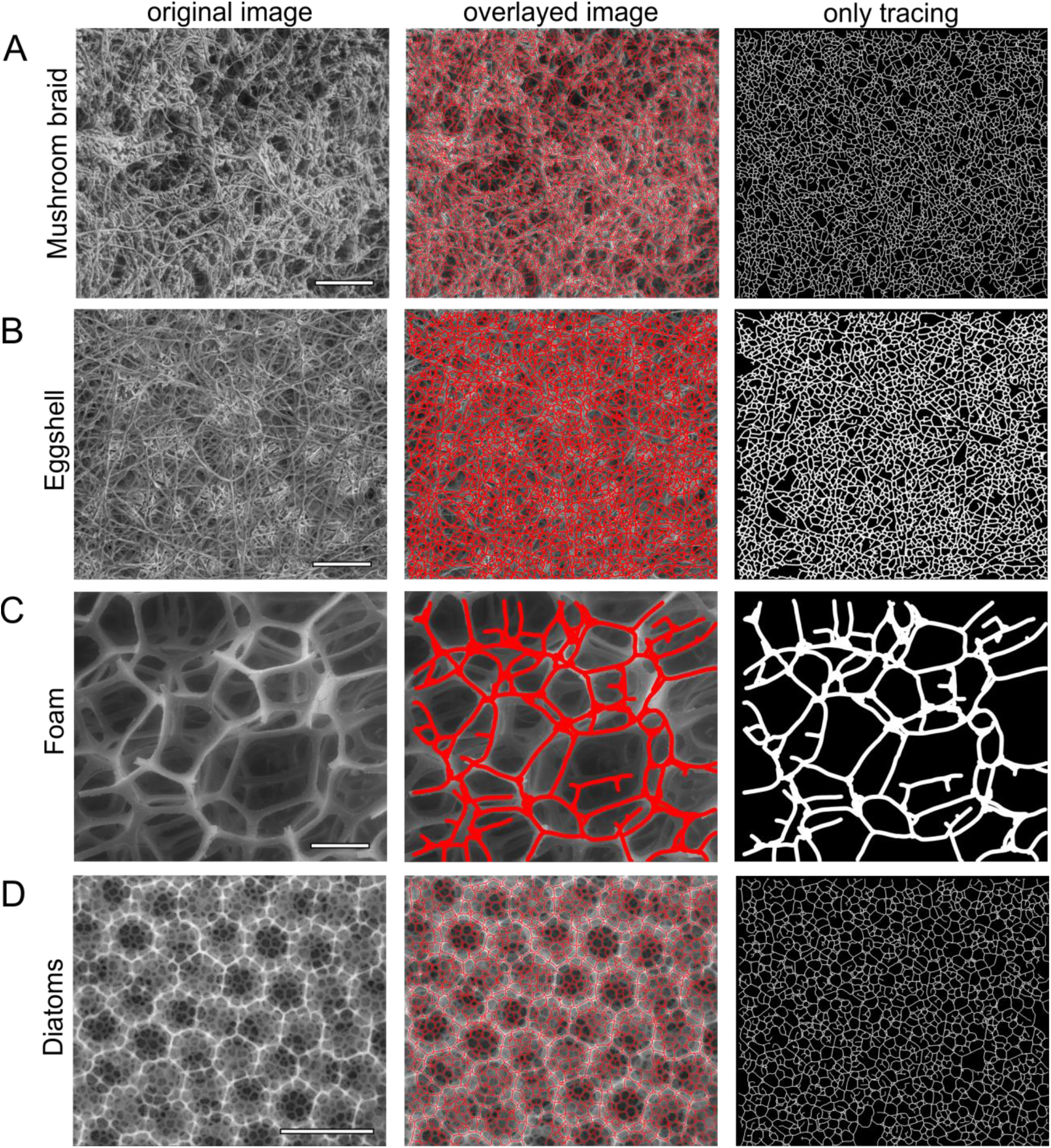
Tracing of various scanning electron microscopy images. Exemplary images and their tracing (as indicated) of mushroom braid (**A**), eggshell (**B**), foam (**C**) and a diatom (**D**). Overlayed image, overlay of original and traced image. Scale bars: 100 μm (A), 50 μm (B), 500 μm (C), 4 μm (D).

**FIGURE S5.**
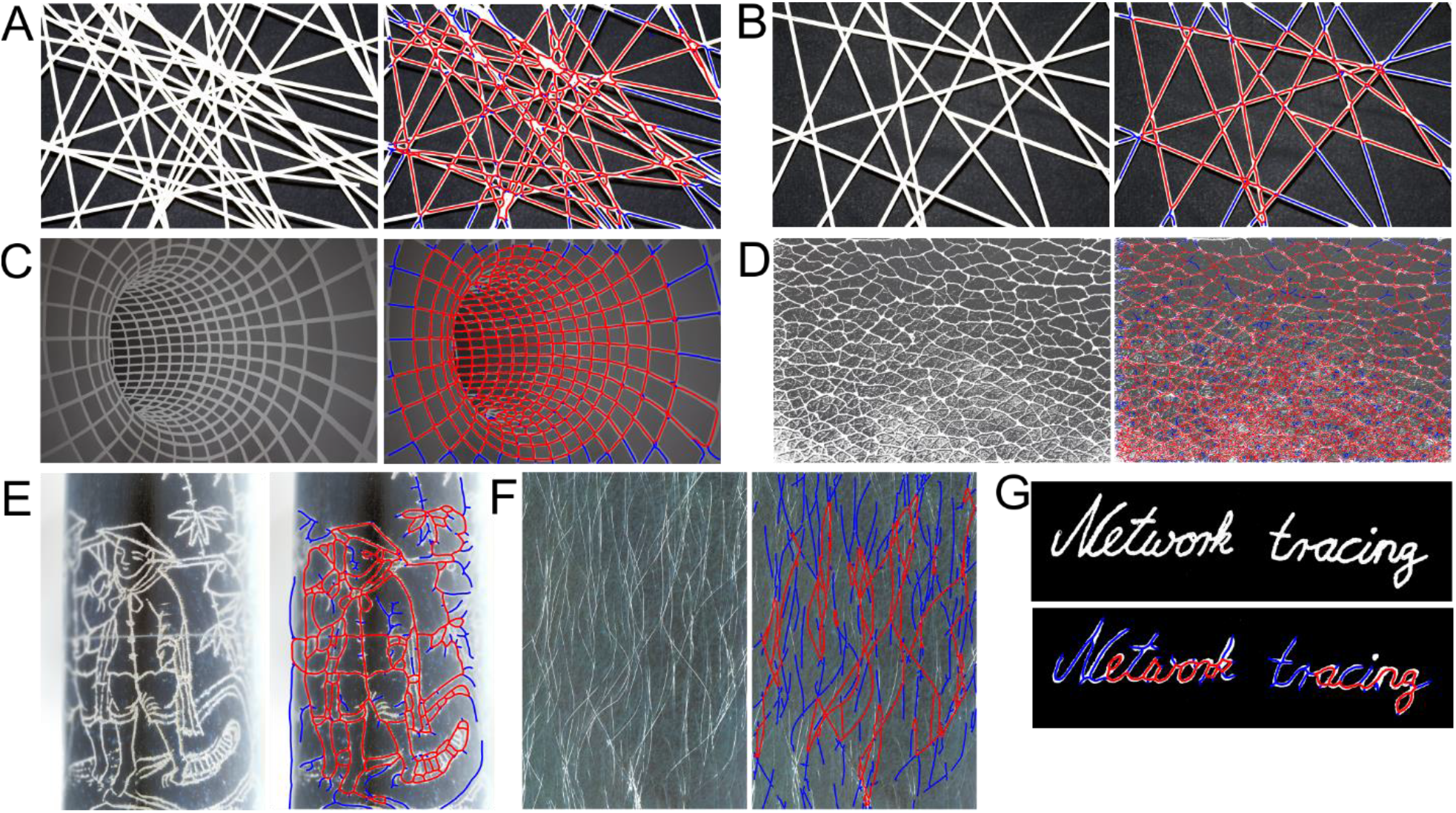
General examples of network tracing using the filament network-tracing algorithm (FiNTA). Exemplary images (left/top) and their tracing (right/bottom), for different kinds of networks using the FiNTA. All of the images were color modified for bright ‘filaments’ and dark backgrounds. The images shown are for: dense (**A**) and porous (**B**) networks of sticks of spaghetti; three-dimensional tunnel painted on a wall (**C**); the structure of leather (**D**); a line drawing of a person (**E**); human leg hairs (**F**); and handwriting (**G**). Red tracing, closed loops; blue tracing, open loops.

## References

1. Xia S, Lim YB, Zhang Z, et al. Nanoscale architecture of the cortical actin cytoskeleton in embryonic stem cells. Cell Rep. J. 2019;28(5):1251–1267 e7.

2. Zemel A, Rehfeldt F, Brown AE, Discher DE, Safran SA. Optimal matrix rigidity for stress fiber polarization in stem cells. Nat Phys. 2010;6(6):468–473.

3. Bovellan M, Romeo Y, Biro M, et al. Cellular control of cortical actin nucleation. Curr Biol. 2014;24(14):1628–1635.

4. Chugh P, Paluch EK. The actin cortex at a glance. J Cell Sci. 2018;131(14)

5. Nasrollahi S, Banerjee S, Qayum B, Banerjee P, Pathak A. Nanoscale matrix topography influences microscale cell motility through adhesions, actin organization, and cell shape. ACS Biomat Sci Engineer. 2017;3(11):2980–2986.

6. Maiuri P, Rupprecht JF, Wieser S, et al. Actin flows mediate a universal coupling between cell speed and cell persistence. Cell. 2015 161(2):374–86.

7. Yamaguchi H, Condeelis J. Regulation of the actin cytoskeleton in cancer cell migration and invasion. Biochim Biophys Acta. 2007;1773(5):642–52.

8. Hall A. The cytoskeleton and cancer. Cancer Metast Rev. 2009;28(1-2):5–14.

9. Cordes A, Witt H, Gallemi-Perez A, et al. Prestress and area compressibility of actin cortices determine the viscoelastic response of living cells. Phys Rev Lett. 2020;125(6):068101.

10. Hecht FM, Rheinlaender J, Schierbaum N, Goldmann WH, Fabry B, Schaffer TE. Imaging viscoelastic properties of live cells by AFM: power-law rheology on the nanoscale. Soft Matter. 2015;11(23):4584–91.

11. Svitkina TM. Actin cell cortex: structure and molecular organization. Trends Cell Biol. 2020;30(7):556–565.

12. Gardel ML, Shin JH, MacKintosh FC, Mahadevan L, Matsudaira P, Weitz DA. Elastic behavior of cross-linked and bundled actin networks. Science. 2004;304(5675):1301–5.

13. Chugh P, Clark AG, Smith MB, et al. Actin cortex architecture regulates cell surface tension. Nat Cell Biol. 2017;19(6):689–697.

14. Chen F, Tillberg PW, Boyden ES. Optical imaging. Expansion microscopy. Science. 2015;347(6221):543–8.

15. Svitkina TM. Platinum replica electron microscopy: imaging the cytoskeleton globally and locally. Int J Biochem Cell Biol. 2017;86:37–41.

16. Chikina AS, Svitkina TM, Alexandrova AY. Time-resolved ultrastructure of the cortical actin cytoskeleton in dynamic membrane blebs. J Cell Biol. 2018.

17. Hotaling NA, Bharti K, Kriel H, Simon CG. DiameterJ: a validated open source nanofiber diameter measurement tool. Biomaterials. 2015;61:327–38.

18. Meijering E, Jacob M, Sarria JC, Steiner P, Hirling H, Unser M. Design and validation of a tool for neurite tracing and analysis in fluorescence microscopy images. Cytometry A. 2004;58(2):167–76.

19. Xu T, Vavylonis D, Tsai FC, et al. SOAX: a software for quantification of 3D biopolymer networks. Sci Rep. 2015;5:9081.

20. Su R, Sun C, Pham TD. Junction detection for linear structures based on Hessian, correlation and shape information. Pattern Recogn. 2012;45(10):3695–3706.

21. Smith MB, Li H, Shen T, Huang X, Yusuf E, Vavylonis D. Segmentation and tracking of cytoskeletal filaments using open active contours. Cytoskeleton (Hoboken). 2010;67(11):693–705.

## References

1. Hahn WC, Counter CM, Lundberg AS, Beijersbergen RL, Brooks MW, Weinberg RA. Creation of human tumour cells with defined genetic elements. Nature. Jul 1999;400(6743):464–8.

2. Hari-Gupta Y, Fili N, dos Santos Á, et al. Nuclear myosin VI regulates the spatial organization of mammalian transcription initiation. bioRxiv. 2020:2020.04.21.053124

3. Chen F, Tillberg PW, Boyden ES. Optical imaging. Expansion microscopy. Science. Jan 30 2015;347(6221):543–8.

4. Chugh P, Clark AG, Smith MB, et al. Actin cortex architecture regulates cell surface tension. Nat Cell Biol. Jun 2017;19(6):689–697.

5. Svitkina TM. Platinum replica electron microscopy: Imaging the cytoskeleton globally and locally. Int J Biochem Cell Biol. May 2017;86:37–41.

